# Risk of Bias Assessment in Preclinical Literature using Natural Language Processing

**DOI:** 10.1101/2021.06.04.447092

**Authors:** Qianying Wang, Jing Liao, Mirella Lapata, Malcolm Macleod

## Abstract

**Objective:** We sought to apply natural language processing to the task of automatic risk of bias assessment in preclinical literature, which could speed the process of systematic review, provide information to guide research improvement activity, and support translation from preclinical to clinical research.

**Materials and Methods:** We use 7,840 full-text publications describing animal experiments with yes/no annotations for five risk of bias items. We implement a series of models including baselines (support vector machine, logistic regression, random forest), neural models (convolutional neural network, recurrent neural network with attention, hierarchical neural network) and models using BERT with two strategies (document chunk pooling and sentence extraction). We tune hyperparameters to obtain the highest F1 scores for each risk of bias item on the validation set and compare evaluation results on the test set to our previous regular expression approach.

**Results:** The F1 scores of best models on test set are 82.0% for random allocation, 81.6% for blinded assessment of outcome, 82.6% for conflict of interests, 91.4% for compliance with animal welfare regulations and 46.6% for reporting animals excluded from analysis. Our models significantly outperform regular expressions for four risk of bias items.

**Conclusion:** For random allocation, blinded assessment of outcome, conflict of interests and animal exclusions, neural models achieve good performance, and for animal welfare regulations, BERT model with sentence extraction strategy works better.

## BACKGROUND

Systematic review is a type of literature review that attempts to collate all empirical evidence relevant to a pre-specified research question. It uses explicit and systematic methods to minimize bias and provide more reliable findings than narrative review [1]. After the collection of research publications which meet prespecified inclusion criteria, a critical step is the reporting of strategies designed to reduce risks of bias (RoB) in the included publications, which is central to the assessment of the reliability of the research findings [2]. The current procedure for risk of bias assessment in literature is that it usually performed separately by two independent investigators, working with an adjudicator to resolve any disagreements. This is both time-consuming and prone to error. As the number of publications describing experimental studies increases rapidly, it has become increasingly difficult for researchers to keep up to date with progress in their field and the findings of systematic reviews are weakened. Therefore, automation tools would accelerate this process and increase reliability. Such tools would also be useful in evaluating the impact of measures designed to improve the quality and completeness of research reporting (NPQIP [3], IICARus [4], MDAR [5]) and in measuring the impact of institutional research improvement activities [6].

Systematic reviewers have advocated the use of automated approaches to assist risk of bias assessment, using human effort and machine automation in mutually reinforcing ways [7]. The development of machine learning and natural language processing (NLP), including neural models and transfer learning, provides opportunities to create robust tools for risk of bias assessment. For clinical trials, RobotReviewer trains support vector machines on 6,610 full texts with pseudo labels derived from 1,400 unique strings of bias domains from the Cochrane Database of Systematic Reviews, which achieves overall accuracy around 71.1% [8]. Zhang et al consider the supported sentence annotations of bias domains as ‘rationales’ and use them to train the convolutional neural networks [9] which improves the performance by 5% compared to baseline models [10]. Millard et al apply logistic regressions on 1,467 full-text clinical reports for sentence and document classification separately and achieves the area under the ROC curve larger than 72% for randomisation sequence generation, allocation concealment and blinding [11]. Menke et al have reported the performance of a proprietary tool ‘SciScore’ [12] which trains the conditional random fields [13] on 250 research articles with manually labelled entity mentions for random allocation and blinding. The training corpus is randomly selected from the PubMed Open Access articles, and the portion of clinical or preclinical publications is not clear.

Compared with clinical trials, animal studies are conducted in relatively small teams, are reported in a different style, have been shown to have lower reporting of strategies to reduce risks of bias [14], and are susceptible to different risks of bias [15]. Hence, separate tools for RoB assessment in preclinical literature are necessary. Bahor et al. have previously reported the use of regular expressions with rule-based string matching to recognize phrases related to RoB reporting in experimental animal studies, which requires many hand-crafted term selections [16]. NLP-based approaches may achieve more robust results in the preclinical literature compared with non-learning algorithms.

## OBJECTIVE

We aim to apply natural language processing to assist automatic risk of bias assessment in the preclinical literature. We implement and compare the performance of eight classification models ranging from baseline approaches to more recent state-of-the-art NLP models for five risk of bias items, and provide recommendations for model selection.

## MATERIALS AND METHODS

We consider the risk of bias assessment as a typical text classification task. A classification model cannot be trained from the plain text directly and we need to convert text information to analysable data. The core concept is to map each document to a matrix consisting of fixed-dimension word vectors or embeddings [17], then train a classification model to map these numeric text representations to the binary RoB label (yes/no). For representation methods, we explore bag-of-words, word2vec [18], doc2vec [19] and embeddings from BERT [20]. For classification models, we implement baseline models (support vector machine, logistic regression, random forest), neural models (convolutional neural network, recurrent neural network with attention, hierarchical neural network) and BERT models using two strategies, which are described in greater detail below. The different approaches are summarized in Figure 1, and training details are given in supplementary materials.

**Figure 1:**
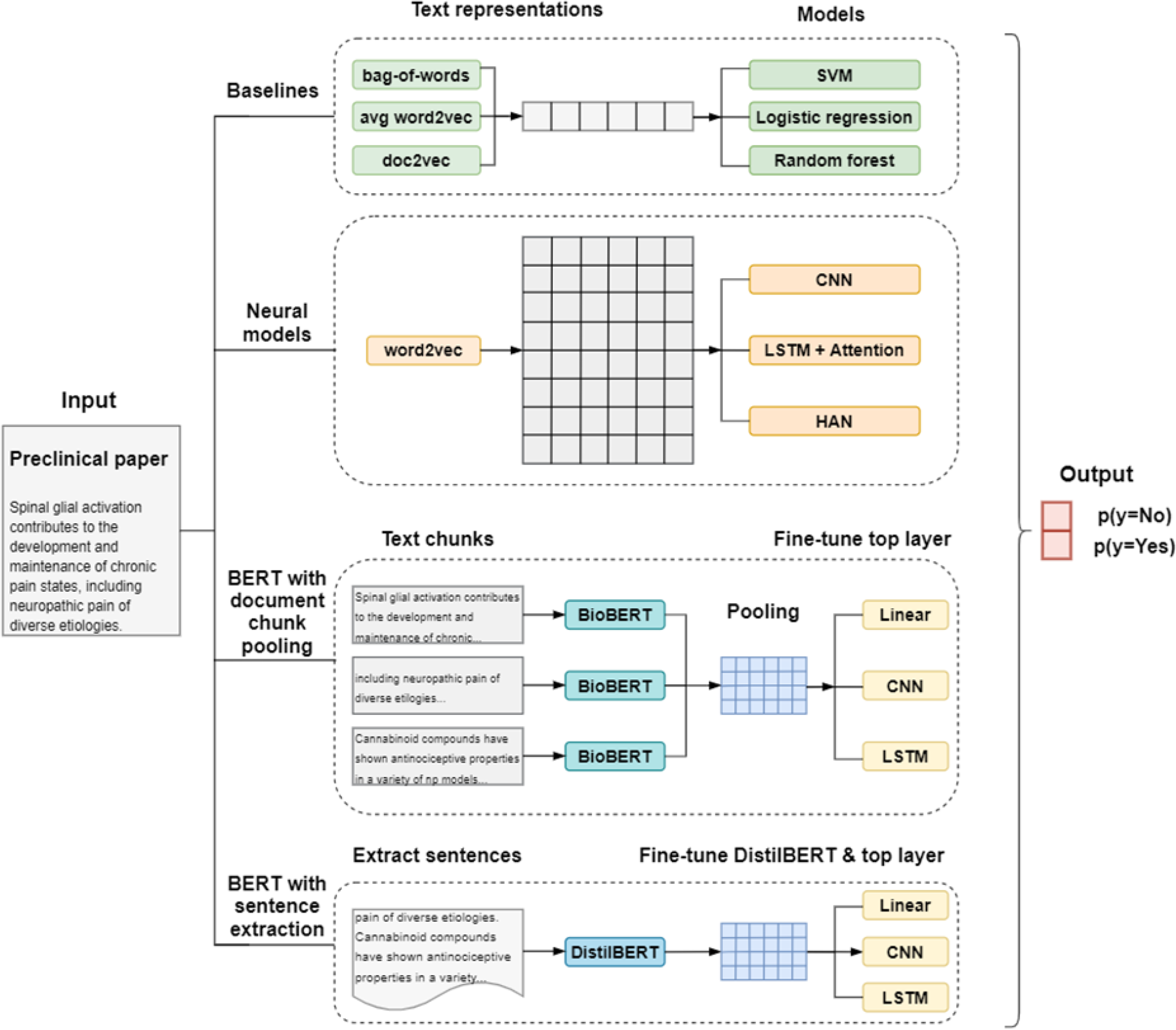
Overall methods of text representations and classification models being tested.

### Dataset

We use a collection of full-text publications which have been annotated for risks of bias [21] in systematic reviews in three research domains (focal ischaemic stroke [22], chemotherapy-induced peripheral neuropathy [23], and psychotic disorders [24]) and in two studies assessing the effectiveness of interventions to improve reporting quality across in vivo research (NPQIP [3] and IICARus [4]). The risk of bias labels are at the document level (1 for reported, 0 for not reported) and each was derived from the annotations of two independent investigators followed by an internal validation process. We consider five risk of bias domains: (1) Random Allocation (RA): animals are randomly allocated to treatment or control groups; (2) Blinded Assessment of Outcome (BAO): group identity is concealed from the scientist measuring the outcome; (3) Compliance with Animal Welfare Regulations (CAWR): researchers report that they complied with relevant animal welfare regulations; (4) Conflict of Interests (CI): authors report any relationship which might be perceived to introduce a potential conflict of interests, or the absence of such a relationship; (5) Animal Exclusions (AE): a statement of whether or not all animals, all data and all outcomes measured are accounted for and presented in the final analysis. Some example sentences indicating the reporting for each risk of bias item are displayed in Table 1.

**Table 1:**
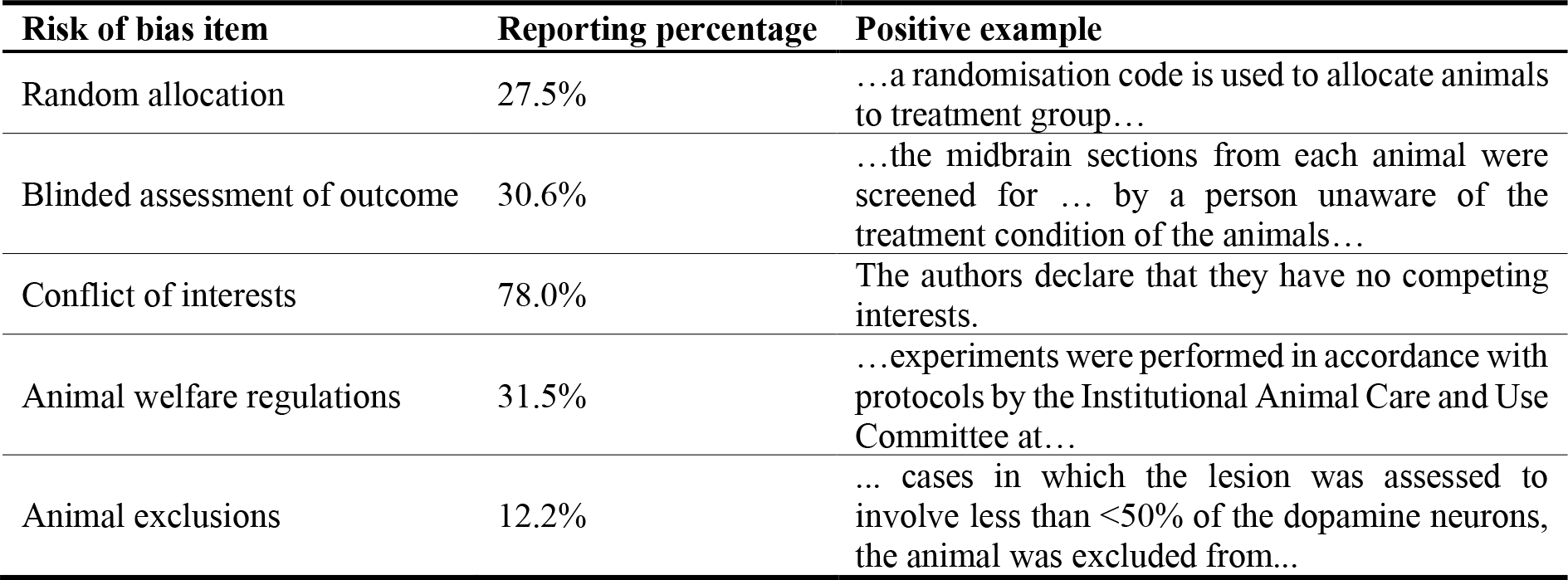
Percentage of papers reporting each risk of bias item, and example sentences from full texts indicating the reporting.

Publications are all in PDF format and we converted them to plain text using Xpdf (https://www.xpdfreader.com). We converted all text to lower case and used regular expressions to remove references, citations, URLs, digits, non-ASCII characters and text which precedes the “Introduction” section, because they are irrelevant to the risk of bias reporting. We used Stanford CoreNLP [25] for word and sentence tokenization. After removing invalid records (for instance where text conversion failed), 7,840 full-text publications had annotations for RA, BAO and AE, and 7,089 had annotations for CAWR and CI. We combined publications from different source projects and randomly allocated them to training (80%), validation (10%) and test (10%) sets. Summary statistics of the dataset are shown in Table 2.

**Table 2:** Data statistics. Samples for random allocation, blinded assessment of outcome and animal exclusions consist of 7,840 records; samples for compliance of animal welfare regulations and conflict of interests consist of 7,089 records.

|  | Samples for RA, BAO, AE |  |  | Samples for CAWR, CI |  |  |
| --- | --- | --- | --- | --- | --- | --- |
|  | Train | Valid | Test | Train | Valid | Test |
| No. documents | 6272 | 784 | 784 | 5671 | 708 | 710 |
| Avg no. tokens per document | 4977 | 5112 | 5077 | 4947 | 5057 | 4964 |
| Avg no. sentences per document | 180 | 186 | 184 | 178 | 182 | 178 |
| Avg no. tokens per sentence | 28 | 28 | 28 | 28 | 28 | 28 |

### Baselines

We explore three text representation methods in baseline models: 1) bag-of-words, 2) word2vec and 3) doc2vec. Bag-of-words (bow) uses word frequency within the document to represent its importance. Considering less important words with high frequency such as ‘the’ and ‘a’, TF-IDF (term frequency-inverse document frequency) weighting is applied, which normalizes the word frequency in a document by multiplying a log-scale of the inverse of the frequency of documents where the word occurred [26]. Word2vec is a neural language model which learns to map words to continuous vectors. It can preserve the semantic relationship among words and can either be generated from the learning process jointly within the classification model or fine-tuned on pre-trained word vectors from other language tasks. As the preclinical literature belongs to the biomedical domain, we use the 200-dimensional word vectors induced on a combination of PubMed and PMC texts with texts extracted from a recent English Wikipedia dump, using the skip-gram model with a window size of 5 [27]. Doc2vec is an unsupervised method which learns to represent a document by a dense vector. There are two approaches for training the dense vector: Distributed Memory (DM) and Distributed Bag-of-Words (DBOW), which is suggested to yield better performance when used together [19].

We explore three baseline classifiers: Support Vector Machine (SVM), logistic regression and random forest. SVM and logistic regression are linear classifiers, which are trained to map the word embeddings to the target RoB label to minimize a hinge loss function and log loss function separately [28]. Random forest is an ensemble-based non-parametric method which combines a number of decision trees trained on various sub-samples [29].

### Neural models

We explore three neural models: Convolutional Neural Network (CNN), a powerful model for text classification [9]; Recurrent Neural Network (RNN) which is good at modelling sequential text data [30]; and Hierarchical Attention Network (HAN) [31] which takes the hierarchical structure among word, sentence and document into consideration. The critical elements in the model architecture are described below and shown in Supplementary Figure 1-3.

#### CNN

We use the classic one-layer CNN [9] for document classification. The main characteristic of CNN is the convolutional layer where multiple filter windows (2D matrices) with different sizes are applied to filter out information. Let *x*[*i*:*j*] denote the matrix extracted from row *i* to row *j* of the document matrix. For one document matrix *x* ∈ ℝ ^*s*×*d*^ and one filter *f* ∈ ℝ ^*h*×*d*^ (where *s* is the document length, *d* is the embedding dimension and *h* is the filter size), the convolution layer sequentially extracts a submatrix which has the same dimension as filter *f* and does the sum operation of the element-wise product between *x*[*i*:*i* − *h* + 1] and *f*. This generates a summarised feature vector *w* ∈ ℝ ^*s* − *h* + 1^ of the document matrix *x* by filter with filter *f* size *h*. For filter size *h*, multiple filters are used to capture different features.

The output vectors from the convolutional layer are then passed through an activation function such as ReLU to add more non-linearity, and a pooling layer, which extracts the maximum value of each vector. A dropout layer, which randomly sets some values in the vectors to zero, is applied to prevent over-fitting. A final linear transformation is applied to map the vector concatenated from the pooling layer into two numeric values, representing separately whether or not the document reported the RoB item.

#### RNN with attention

Recurrent neural network (RNN) is a type of neural network which builds connections over time steps [32]. In the hidden layer, by combining the weighted hidden representations from the previous word and the next word (if it is applied bidirectionally) through a Tanh operation, a basic recurrent neural structure can retain information in the text from both directions. RNN can handle any-length texts and but if the sequence is very long, it is difficult to keep the information from very earlier steps to later steps because of the exploding or vanishing gradient problem [33]. Two variants of RNN, LSTM [30] and GRU [34] are designed to solve this long-term dependencies problem, which uses multiple gates (forget gate, input gate and output gate in LSTM; reset gate, update gate and output gate in GRU) for each word embedding to control the information we need to flow straight, forget, store and update to the next step.

In the general RNN structure, the output from the hidden layer is obtained by simply taking the hidden state of the last RNN cell, which loses some information from other RNN cells; or averaging hidden states of all RNN cells, which treats words at different positions equally. However, the same word may play a different role in the decision of the classification when it occurs in different sentences or contexts. A global context matrix (∈ ℝ^*s*×*h*^) is created to learn the importance of each word in the document (similar to the attention mechanisim described in HAN). The attention module is then added to learn and emphasize the word contributions to the entire document sequence [35].

#### HAN

Words contribute differently to an individual sentence and sentences contribute differently to the whole document. HAN is proposed to imitate this hierarchical structure of documents, having two levels of attention modules applied at word-level and sentence-level [31]. After the RNN hidden layer, in the word-level attention module, the hidden representations of each word in a sentence are multiplied by a local word context vector, which is trained to learn the importance of each word in the sentence. The representation vector of each sentence is then summarised from those weighted word representations. Similarly, in sentence-level attention, the hidden representations of each sentence in the document are multiplied by a global sentence context vector, which is trained to learn the importance of each sentence in the document. Then a document representation vector is obtained from those new weighted sentence representations. After an activation function and a linear transformation, we then output the probability for RoB items. With the hierarchical structure, HAN can generate ranking scores for sentences, which can be used to extract the most relevant sentences and provided to users to allow them to make a judgment on the veracity of the machine decision.

### BERT models

One limitation of word embeddings like word2vec is that the representation vector of a given word is fixed and independent, regardless of context. Contextualized representation models like BERT [20] address this issue. BERT extracts the contextualized embeddings by training a deep bidirectional encoder from transformers [36] on the BooksCorpus and English Wikipedia. The Transformer structure mainly consists of identical blocks, and each block contains sub-modules based on multi-head self-attention and a feed-forward neural network. It dispenses with recurrence and convolutions, and achieves state-of-the-art performance on many natural language processing tasks [36]. The pre-trained BERT can be fine-tuned with a simple additional output layer for downstream tasks. BERT uses WordPiece with a 30,000 token vocabulary for tokenization, which handles rare words better than the ‘pure’ word embeddings and more efficiently than character embeddings [37].

Previous work shows that the domain corpus used for pre-training affects the performance of the downstream task [38]. Since our task is conducted on preclinical texts, we use the pre-trained weights from BioBERT to initialize the model, which applies the same architecture as BERT and is pre-trained on combinations of text corpora including BookCorpus, English Wikipedia, PubMed abstracts and PubMed Central full-text articles [39].

One drawback of BERT is that it can only accept embeddings of maximum 512 tokens as input, which limits the usage for tasks with long documents. There are other transformer models designed for long documents, such as Longformer [40] which can process a maximum of 4096 tokens. However, this is still computationally expensive, and our full-text publications contain 5000 tokens on average. To solve this issue, we propose two strategies.

#### BERT with Document Chunk Pooling (BERT-DCP)

We split documents into text chunks, apply BioBERT to each chunk, and pool the hidden states from different chunks using multiple strategies. This is similar to the structure applied in the classification of clinical notes for patient smoking status [41], with some modifications as shown in Supplementary Figure 4. After the WordPiece tokenization, considering a document with s tokens, the document is split into m=⌈s/510⌉ chunks (excluding the first token [CLS] indicating classification and separation token [SEP] for sentence segmentation). The input representation of the document is X ∈ ℝ^*m*× 512 × *h*^, where h is the hidden dimension throughout the embedding layer and encoder layers in BioBERT. Instead of taking the hidden states from the last encoder layer, we perform the average pooling operation over several encoder layers to obtain the output. We summarize across tokens within each chunk with five different options: 1) max pooling, 2) average pooling, 3) concatenate output from max pooling and average pooling, 4) use hidden states of the [CLS] token, 5) concatenate hidden states of all tokens. After two pooling layers, we explore three head layers (linear/convolutional/LSTM) for the downstream classification task. The convolution and LSTM head use the same architecture as described previously. Unlike convolution or LSTM head, the linear head cannot handle sequences of different lengths, so we add another pooling layer to obtain the fixed-dimension output. The pooling methods use the same options applied in the second pooling layer, with the exclusion of ‘concatenate hidden states of all tokens’, which does not generate a fixed-dimension output.

#### BERT with Sentence Extraction (BERT-SE)

Instead of using the full-text document as input, we extract the most relevant sentences to the risk of bias description. We first use scispaCy [42] to split a document into sentences, and then apply SentenceTransformers [43] to obtain a vector for each individual sentence. We also feed a description sentence of each RoB item (see descriptions in Dataset) to the SentenceTransformers and obtain the corresponding representative vectors. For each individual document, we calculate the cosine similarity score between each sentence vector and the vector of the RoB description sentence. We take the first *k* sentences with the highest similarity scores, i.e. the most *k* relevant sentences, to form a new shorter passage. We then fine-tune the DistilBERT [44] model (a smaller, faster and lighter version of BERT), with a linear/convolution/LSTM head on the new passage, to generate the probabilities of RoB reporting. The sentence extraction process is unsupervised and is independent of the actual training process.

## RESULTS

The results of eight models from three categories (baselines, neural models, and models using BERT with two strategies) on the validation set are shown in Table 3. For baseline models, all items achieve F1 score over 48% and particularly, models for compliance with animal welfare regulations show good performance, with F1 around 90%. For the selection of text representation methods, from our experiments, bag-of-words is not robust and prone to over-fitting. Doc2vec gives the best results across all items, because the training sample texts for doc2vec are closer to the preclinical domain, while the pre-trained word2vec vectors are induced from the more general biomedical corpus. For model selection, logistic regression achieves the best performance for random allocation to treatment or control, blinded assessment of outcome and conflict of interests; while for compliance with animal welfare regulations and animal exclusions, SVM performs better.

**Table 3:** Performance of best model in three categories (baseline, neural model, and BERT models with two strategies) for risk of bias items on the validation set.

| **RoB item** | **Model** | **F1** | **Recall** | **Precision** |
| --- | --- | --- | --- | --- |
| **Random allocation** | SVM | 51.9 | 72.2 | 40.5 |
|  | LogReg | 55.3 | 73.7 | 44.3 |
|  | RF | 67.2 | 79.9 | 58.1 |
|  | CNN | 86.4 | 93.2 | 81.8 |
|  | **RNN+Attn** | **87.2** | 92.4 | 83.7 |
|  | HAN | 86.2 | 91.3 | 83.1 |
|  | BERT-DCP | 85.4 | 92.7 | 80.1 |
|  | BERT-SE | 80.6 | 82.0 | 82.0 |
| **Blinded assessment of outcome** | SVM | 59.3 | 67.8 | 52.7 |
|  | LogReg | 60.0 | 69.1 | 53.0 |
|  | RF | 57.8 | 68.3 | 50.2 |
|  | CNN | 82.4 | 88.5 | 77.8 |
|  | RNN+Attn | 83.0 | 91.1 | 77.2 |
|  | HAN | 81.3 | 86.4 | 77.5 |
|  | **BERT-DCP** | **83.1** | 91.8 | 77.0 |
|  | BERT-SE | 79.9 | 84.7 | 79.8 |
| **Conflict of interests** | SVM | 67.1 | 79.7 | 57.9 |
|  | LogReg | 68.8 | 76.1 | 62.8 |
|  | RF | 65.1 | 68.5 | 61.9 |
|  | CNN | **84.5** | 86.8 | 84.1 |
|  | RNN+Attn | 82.9 | 85.4 | 82.0 |
|  | HAN | 83.2 | 84.7 | 82.8 |
|  | BERT-DCP | 79.5 | 84.6 | 76.8 |
|  | BERT-SE | 64.0 | 64.3 | 70.9 |
| **Compliance of animal welfare regulations** | SVM | 90.1 | 96.3 | 84.6 |
|  | LogReg | 87.6 | 85.4 | 89.9 |
|  | RF | 88.8 | 89.7 | 88.0 |
|  | CNN | 86.9 | 83.3 | 92.4 |
|  | RNN+Attn | 76.3 | 77.6 | 78.3 |
|  | HAN | 79.3 | 77.9 | 84.5 |
|  | BERT-DCP | 93.8 | 92.1 | 95.8 |
|  | **BERT-SE** | **94.0** | 94.6 | 93.8 |
| **Animal exclusions** | SVM | 39.0 | 64.3 | 28.0 |
|  | LogReg | 41.4 | 62.5 | 31.0 |
|  | RF | 48.8 | 44.6 | 53.8 |
|  | CNN | **60.2** | 73.6 | 54.2 |
|  | RNN+Attn | 58.0 | 68.3 | 54.3 |
|  | HAN | 53.4 | 58.4 | 54.0 |
|  | BERT-DCP | 56.2 | 77.0 | 46.8 |
|  | BERT-SE | 36.7 | 56.6 | 29.2 |

Neural models are more robust to hyperparameter tuning than baseline models in our experience. For random allocation, blinded assessment of outcome and conflict of interests, neural models improve the performance by 14% to 30% over baseline models, and the difference of results among three neural models are not obvious. For compliance with animal welfare regulations, neural models do not show advantages over baselines, with performance reduction ranging from 4% to 14%. For animal exclusions, weight balancing strategy and under-sampling do not reduce the effect of data imbalance issue, and the training process is prone to over-fitting.

Models using BERT with the two strategies described do not outperform neural models, except item CAWR, which has 3%∼4% improvement. This is reasonable because in the document chunk pooling strategy, we do not take any advantages of BERT structure by freezing all the encoder layers, and multiple pooling strategies help little to address this limitation; in the sentence extraction strategy, although we can fine-tune DistilBERT, we still lose some information by using shorter texts extracted from full publications. We have not been able to evaluate the performance of sentence extraction modules, which requires further sentence-level annotations.

With the best model and its optimal setting for each risk of bias item, we evaluate and compare the performance with the regular expression approach on the test set. Note that we select RNN with attention as the optimal model for blinded assessment of outcome rather than BERT with document chunk pooling strategy, considering the negligible improvement (0.1%) and complexity of pre-processing by the latter approach. From Table 4, our NLP models improve performance by between 13% and 36% for four RoB items tested, and these improvements are significant with p < 0.05 according to McNemar’s test [45].

**Table 4:** Performance of the best NLP model and regular expression approach for each risk of bias item on the test set. A regular expression approach has not been developed for animal exclusions.

| RoB item | Model/Approach | F1 | Recall | Precision |
| --- | --- | --- | --- | --- |
| Random allocation | RNN+Attn | **82.0** | 86.8 | 79.5 |
|  | Regular expression | 68.8 | 96.4 | 53.6 |
| Blinded assessment of outcome | RNN+Attn | **81.6** | 87.8 | 78.2 |
|  | Regular expression | 68.3 | 59.8 | 79.6 |
| Conflict of interests | CNN | **82.7** | 80.6 | 86.2 |
|  | Regular expression | 48.7 | 33.8 | 87.1 |
| Compliance with animal welfare regulation | BERT-SE | **91.5** | 91.4 | 92.0 |
|  | Regular expression | 55.2 | 40.9 | 85.2 |
| Animal exclusions | CNN | 46.6 | 56.5 | 45.0 |
|  | Regular expression | -- | -- | -- |

Supplementary Table 1 demonstrates the prediction and sentence extraction function of our models on an example paper which reports RA, BAO and AE, but does not report CI and CAWR. Unlike the previous rule-based approaches which output yes/no label only, our models can be used to extract the most relevant sentences from full text, which can enhance the judgment from the prediction probabilities, or provide signals whether users need to re-check the full texts. In Supplementary Table 1, sentences extracted for RA, BAO and AE indicate the clear relation with the items and positive evidence for the prediction probabilities, while sentences extracted for CI and CAWR do not show any relation with the items, which proves the predictions in a different direction.

## DISCUSSION

We have shown that different models are optimal for the detection of reporting of different risks of bias. CNN is the best choice for conflict of interests and RNN with attention works well for random allocation to groups and blinded assessment of outcome. For compliance with animal welfare regulations, models using BERT with sentence extraction strategy achieve the best performance. For animal exclusions, CNN achieves the best performance on the validation set, but no approach provides reliable performance on the test set. Compared with the previous regular expression approach, the F1 scores for four risk of bias items are between 13% and 36% higher, indicating a substantial improvement. The sentence extraction function can provide potentially relevant sentences as clues for users making judgment.

We analyse all positive samples and use RNN with attention module to output attention scores for tokens in each individual paper, thus we can extract the most important words in the decision of classification task. The five most important words are {“randomly”, “induced”, “supported”, “randomized”, “increase”} for random allocation, {“blind”, “by”, “observer”, “experimenter”, “investigator”} for blinded assessment of outcome, {“interest”, “of”, “no”, “authors”, “statement”} for conflict of interests, {“animal”, “care”, “procedures”, “figure”, “committee”} for animal welfare regulations, and {“excluded”, “were”, “from”, “included”, “died”} for animal exclusions (Figure 2). This may help future rule-based approaches development.

**Figure 2:**
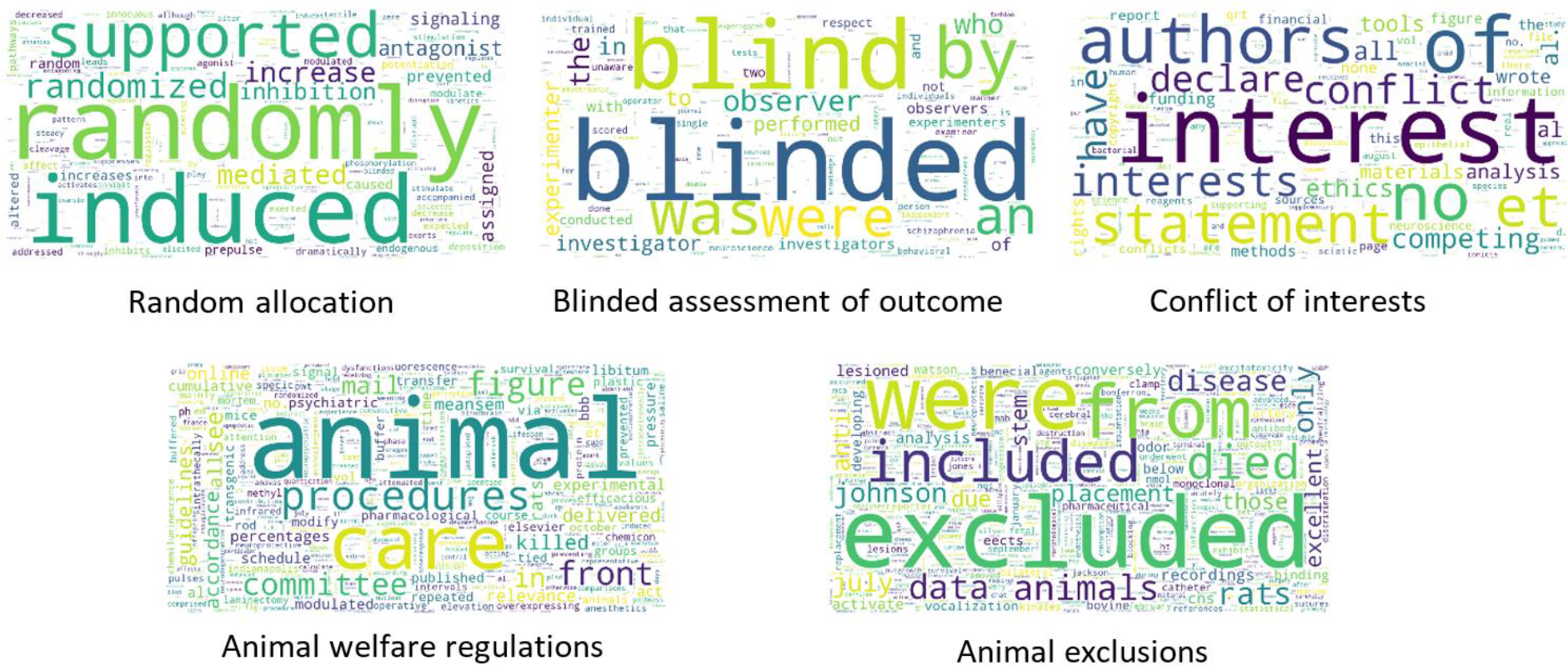
Most important words in the decision of classification for each risk of bias item, based on the average attention scores from RNN output over all positive samples.

Among the incorrect records, our models are more likely to conclude that papers report random allocation, blinded assessment of outcome and animal exclusions (false positive greater than false negative), and less likely to predict that papers report conflict of interests and animal welfare regulation (false negative greater than false positive, Figure 3). To analyse sources of error we randomly selected 10 incorrect records for each item from the test set. Our models did not recognise phrases like ‘unaware’ for blinded assessment but considered that ‘animals are randomly selected for testing’ indicated random allocation to the experimental group. It may be that most records in our training set describe random allocation based on the presence of the word ‘random’ and blinded assessment based on the word ‘blind’, and that our training corpus did not have sufficient examples of alternative valid descriptions for these to be learned. We also found two records where a conflict of interests was given before the ‘Introduction’ section or after the ‘Reference’ section, where we had removed the relevant text in the text processing stage.

**Figure 3:**
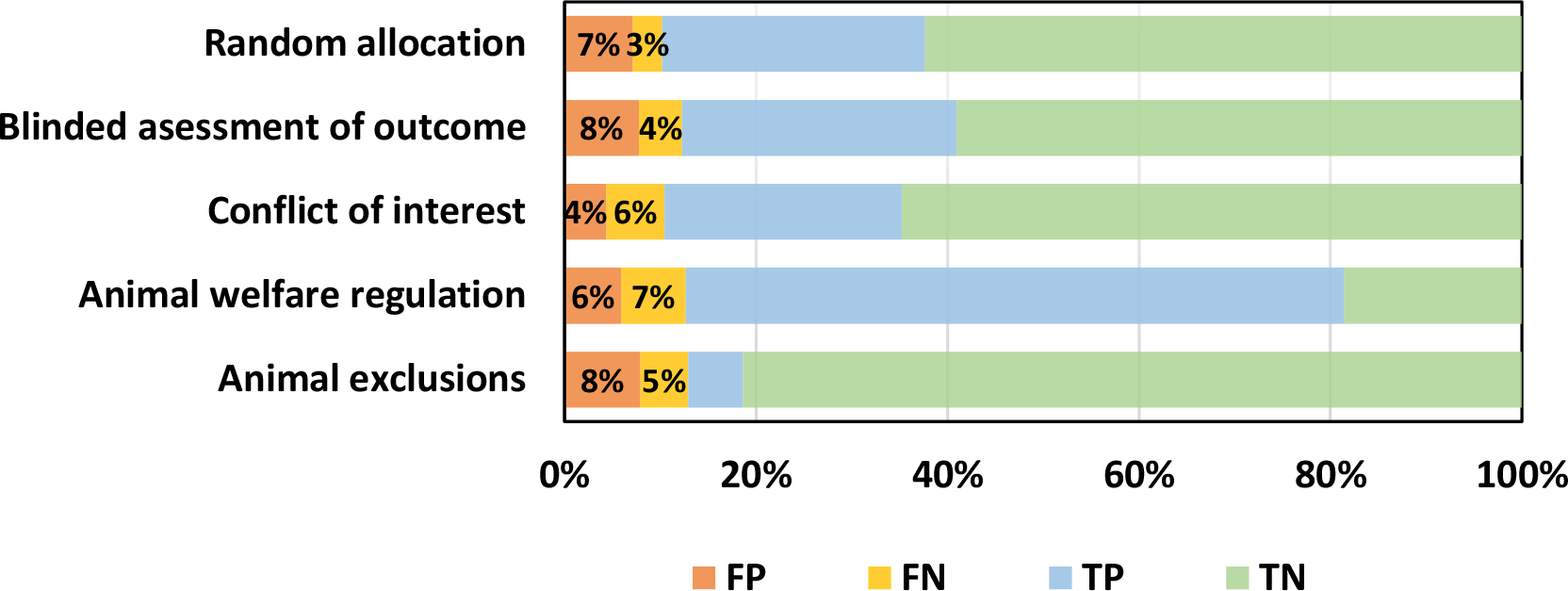
Percentages of false positive, false negative, true positive and true negative of each optimal model for the corresponding risk of bias item on the test set.

The code for predicting probabilities of risk of bias reporting in preclinical full texts is available at https://github.com/qianyingw/rob-pome. The levels of performance achieved make these tools suitable for research improvement activity where several hundred publications are to be evaluated. For instance, for random allocation in a corpus of 1000 manuscripts, this approach would estimate prevalence within 3% of the true value and for 100 publications, within 10% of the true value (see calculations at https://github.com/camaradesuk/confidence_intervals_simulation). Given that the changes sought in research improvement activities are at least of this magnitude, we consider the performance of these tools in determining the reporting of risk of bias items to be such that they are suitable for deployment in a research improvement context. Similarly, they are suitable for the evaluation of risk of bias in large corpuses such as collections in the preclinical systematic reviews. However, they are not yet at the level where they are appropriate for the evaluation of individual publications.

Our work has several limitations. First, our training dataset includes publications drawn from three datasets focusing on specific disease models (focal ischaemic stroke, chemotherapy-induced peripheral neuropathy, psychotic disorders), as well as two datasets from unselected preclinical studies published in PLOS One and Nature. This may influence the generalizability of our findings. Second, PDF to text conversion loses document structure and we cannot identify the main sections of publications. This introduces some noise (for instance text from figures and tables) to our training corpus. Tools like GROBID (https://github.com/kermitt2/grobid) can convert PDFs to structured XML but it highly depends on the quality of PDF, and in our experience it does not work well for some preclinical publications. However, enhanced approaches to PDF conversion, and increased availability of publications in XML format, means that this approach may become feasible in the future.

In future work we will seek to improve performance further, using datasets involving more journals and a wider range of preclinical experiments (both disease modelling and mechanistic studies), and will exploit diseases and texts from structured PubMed XMLs, which may yield better performance. We will continue improving the attribution of animal exclusions to achieve more reliable performance and we will develop approaches for other risk of bias items including sample size calculation and allocation concealment. We will also develop a user-friendly function embedded in the preclinical systematic review facility SyRF (http://syrf.org.uk/) and a standalone API, enabling usage to others.

## CONCLUSION

We explore multiple text classification models, from baselines to recent NLP techniques and demonstrate the advantages of neural models and BERT models for risk of bias assessment in preclinical literature. BERT models work well for animal welfare regulations, while CNN/RNN achieves better performance for random allocation, blinded assessment of outcome, conflict of interests and animal exclusions. We encourage the use of NLP techniques to assist risk of bias assessment and reduce workflow for the preclinical systematic review. If computational limitations require the implementation of a single tool, we recommend neural models like CNNs. The performance of these tools is such that they could be deployed in automated approaches to monitor risks of bias reporting as part of institutional research improvement activities.

## ACKNOWLEDGMENTS

We would like to thank Sarah McCann, Gillian Currie, Zsanett Bahor and Kaitlyn Hair for providing the datasets. This work is jointly funded by China Scholarships Council, John Climax UK Reproducibility Network PhD studentship, and the University of Edinburgh.

