## Supplementary Material for "Risk of Bias Assessment in Preclinical Literature using Natural Language Processing"

### A. Training and hyperparameter details

In all the experiments, we use F1 score (the harmonic mean of recall and precision) as the main metric to select the optimal parameters for each model in the validation set. We also report recall and precision, measuring how well the model is able to detect papers which truly reported RoB, and how reliable it is when the model concludes that papers reported RoB. Models are implemented using Python libraries including scikit-learn [1], PyTorch [2] and Huggingface Transformers [3]. The experiments of baseline models are conducted using a CPU with 16 cores and other experiments are conducted on a GTX 1080 GPU.

#### A.1 Baseline models

We tune minimum document frequency (we ignored words with a document frequency lower than this threshold) and maximum vocabulary size (we only considered the most abundant words ordered by term frequency when building the corpus) for all three text representation methods. For bag-of-words, we use l2 normalization for TF-IDF weighting; we tune word n-gram size, with or without TF-IDF weighting. For word2vec, we average all the word vectors to generate a new feature vector for each document because all three baseline models require a single dimension vector as input. For doc2vec, we set the initial learning rate to 0.01 and reduce it linearly to 0.001 during 20 training epochs; and we draw 5 'noise words' (negative sampling). We average the context vectors to generate the document vector. We tune the dimension of feature vectors generated from the model and training algorithm (DM, DBOW, or combination of DM and DBOW).

For classification models, we assign class weight to solve data imbalance issue, calculated as the number of samples/(number of classes * number of samples within the class). For SVM, we tune the learning rate. 10% of the training samples are used for early stopping to terminate training. For logistic regression, we tune the inversed value of regularization strength. For random forest, we set the maximum depth of individual trees to 2. We tune the number of trees in the forest, and the maximum number of features considered in the splitting process, which is selected from the total number of features, $sqrt$(total number of features) or $log_2$(total number of features).

#### A.2 Neural models

For text representations, we train the pre-trained biomedical word vectors jointly within our neural network models. For all three models, we set the batch size to 32 and dropout rate to 0.5. When building the vocabulary, we consider the 5000 most frequent tokens (words) as features from the training corpus and we exclude tokens occurring in fewer than 10 documents. We explore under-sampling strategies for animal exclusions because it has very imbalanced classes (positive 12.2% / negative 87.8%). We use different early stopping criteria for each RoB item. Parameters are trained to minimize the cross-entropy loss using Adam algorithm [4] with an initial learning rate of 1e-4.

**CNN.** For a starting configuration, we set the filter size to {3, 4, 5} and apply 100 filters for each filter size. Documents are padded by zero or cut when their token length is less or more than 5000. With this

[1] Centre for Clinical Brain Sciences, University of Edinburgh, UK
[2] School of Informatics, University of Edinburgh, UK

setting unchanged, we tune the maximum document length, filter sizes, and number of filters for each filter size, maximum number of features, assigning class weight or not, applying batch normalization layer or not, and freeze embedding or not. Based on some initial experiments for CNN model convergence, we set number of training epochs to 20 for RA, BAO and CI, and 40 for CAWR and AE.

**RNN with Attention.** We first use LSTM module in the one-layer bidirectional RNN structure and set hidden states dimension to 50. We set the maximum document length to 10000. We then explore the effect of different structure of recurrent cell, i.e. LSTM and GRU. We tune the dimension of hidden states, number of hidden layers, maximum document length, maximum number of features, applying bidirectional structure or not, assigning class weight or not, applying batch normalization layer or not, and freeze embedding or not. For the context vector in attention mechanism, we use Kaiming initialization [5] to take care of variance of weights which helps prevent the gradients grow or shrink. Based on some initial experiments for RNN model convergence, we set number of training epochs to 40 for compliance with animal welfare regulations and 20 for the other four items.

**HAN.** For a starting configuration, documents are padded or cut when its number of sentences is less or more than 500, and each sentence is padded or cut if the number of words is less or more than 100. We set dimension of word hidden states and sentence hidden states to 50. We tune the maximum number of sentences, maximum number of words in each sentence, dimension of word hidden states and sentence hidden states, maximum number of features, assigning class weight or not and applying batch normalization or not. We use Kaiming to initialize word context vector and sentence context vector. From some initial experiments, we set the number of training epochs to 20 for RA, BAO and CI, and 30 for the other two items.

#### A.3 BERT models

In the document chunk pooling strategy, each document is tokenized by WordPiece and then split into chunks with 512 tokens. We set the maximum number of chunks to 20 due to the memory issue. Therefore, the model can process maximum 10,200 (512x20) tokens from each document. The last chunk with fewer than 512 is padded. We use the default configuration of the BERT-base module [6]. For output of each BERT chunk, we perform average pooling over last 10 encoder layers from the BERT output in the first pooling layer; we concatenate hidden states of all the tokens in every chunk (no pooling) in the second pooling layer. Due to the memory issue, we freeze all the BERT layers and only fine-tune the linear/convolution/LSTM head for the classification task. For BERT with linear head, we use the average pooling method for the third pooling layer; for BERT with convolution head, we set three filter sizes to {3,4,5} and apply 100 filters for each filter size; for BERT with LSTM head, we use one-layer unidirectional LSTM, and the hidden dimension is set to 768.

We apply gradient clipping with a threshold norm of 0.1 to rescale gradients [7] and gradient accumulation every 16 steps (mini-batches) to reduce memory consumption. Parameters are trained to minimize the cross-entropy loss using AdamW algorithm [8]. We use the slanted triangular learning rate scheduler [9] with maximum learning rate 1e-3.

With those settings, we run each model for 30 epochs. Models with linear head or LSTM head do not show robust performance, with the highest F1 score of around 40%. Considering the running cost and the performance, we continue experiments for models with convolution head only. We tune three hyperparameters for item RA first: the number of encoder layers being averaged in the first pooling layer, and the pooling method in the second pooling layer as described in Materials and Methods (BERT models). Our experiments for item RA indicate that varying the number of encoder layers averaged in the first pooling layer has minimal effect on performance. In the second pooling layer, pooling the output over tokens within each chunk with different methods reduces performance by 40%, compared with concatenating the hidden representations of all the tokens within each chunk. We also explored unfreezing and fine-tuning the last encoder layer in BERT, but this reduces performance by 30%. As the performance does not indicate much improvement, also considering the running cost, we did not continue tuning parameters for the other four RoB items, rather choosing the default setting.

In the sentence extraction strategy, we do not need to freeze any encoder layer and can fine-tune them together with the linear/convolution/LSTM head, because the texts used for training are much shorter than full texts in document chunk pooling strategy and DistilBERT has less memory consumption compared with BERT. For the sentence retrieval module, we tune the number of sentences extracted. We set batch size to 16 and the maximum learning rate in the slanted triangular learning rate scheduler to 5e-2. Other settings are same as that of document chunk pooling strategy.

### B. Tables and figures

Supplementary Table 1: An example of model predication and relevant sentence extraction for risk of bias items on a full-text paper (PMCID: PMC6579011). Prediction probabilities are generated from the optimal model of each item, and most relevant sentences are extracted by HAN.

| Risk of bias item | True | Prediction | High-scored sentences |
|---|---|---|---|
| Random allocation | Yes | 99.97% | In the last 5 min of this habituation period, three 5 sec, 56 dB, substartle-threshold white noise tones were presented randomly by computer. |
| Blinded assessment of outcome | Yes | 99.99% | Video records of 11 randomly selected animals were recoded by an observer blind to the experimental conditions. |
| Conflict of interests | No | 0.32% | Schematic depictions of the regions dissected for neurochemical analysis are presented in Figure 2. |
| Animal welfare regulations | No | 3.68% | Role of the Amygdala in the Coordination of Behavioral, Neuroendocrine, and Prefrontal Cortical Monoamine Responses to Psychological Stress in the Rat. |
| Animal exclusions | Yes | 99.99% | In 8 of the original 26 lesioned animals in the pretraining experiment, the lesions were judged incomplete by the criteria above and were excluded from the data analyses. |

Supplementary Figure 1: An example of CNN for document classification. A document with 8 words is mapped by a 5-dimension word embedding. Two filter sizes [3,4] are used, and each of them has two filters.

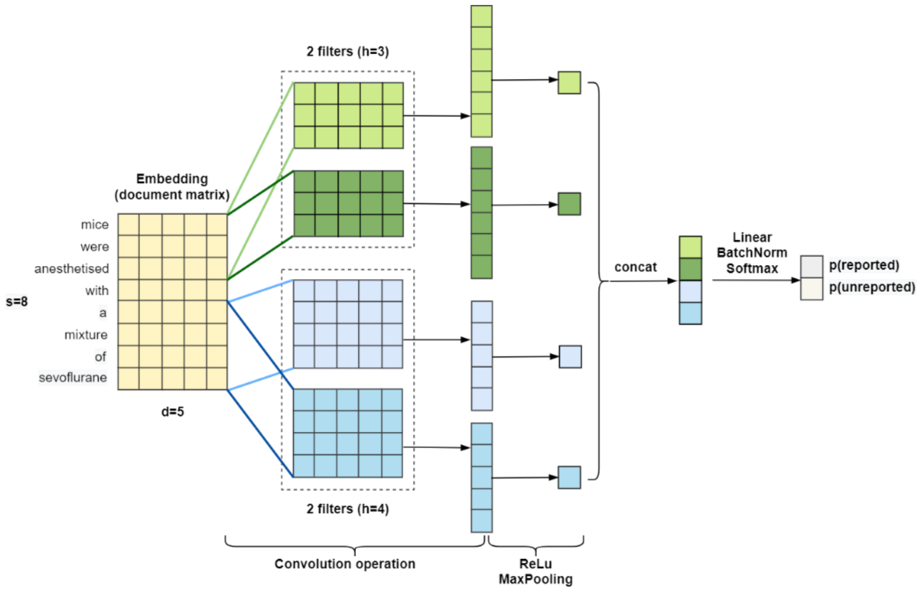

Supplementary Figure 2: One-layer bidirectional RNN with attention mechanism.

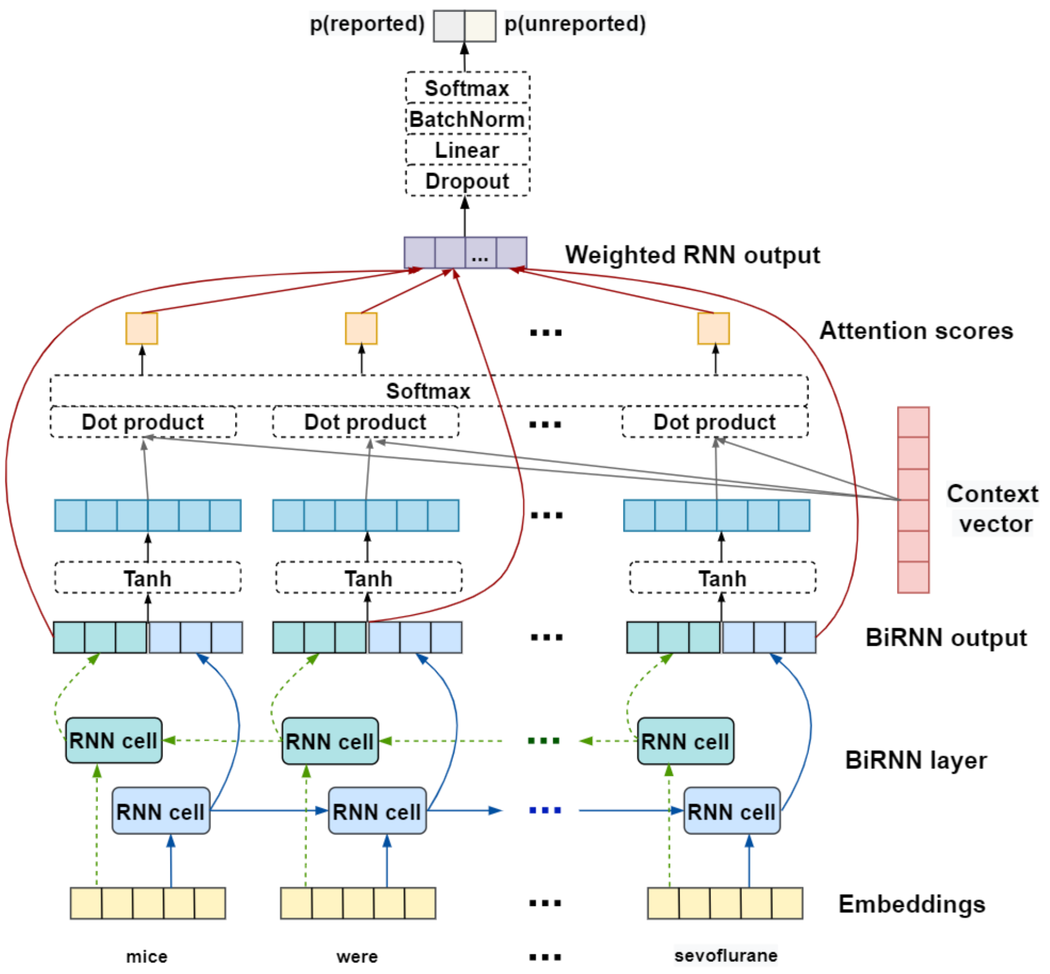

Supplementary Figure 3: The architecture of hierarchical attention network.

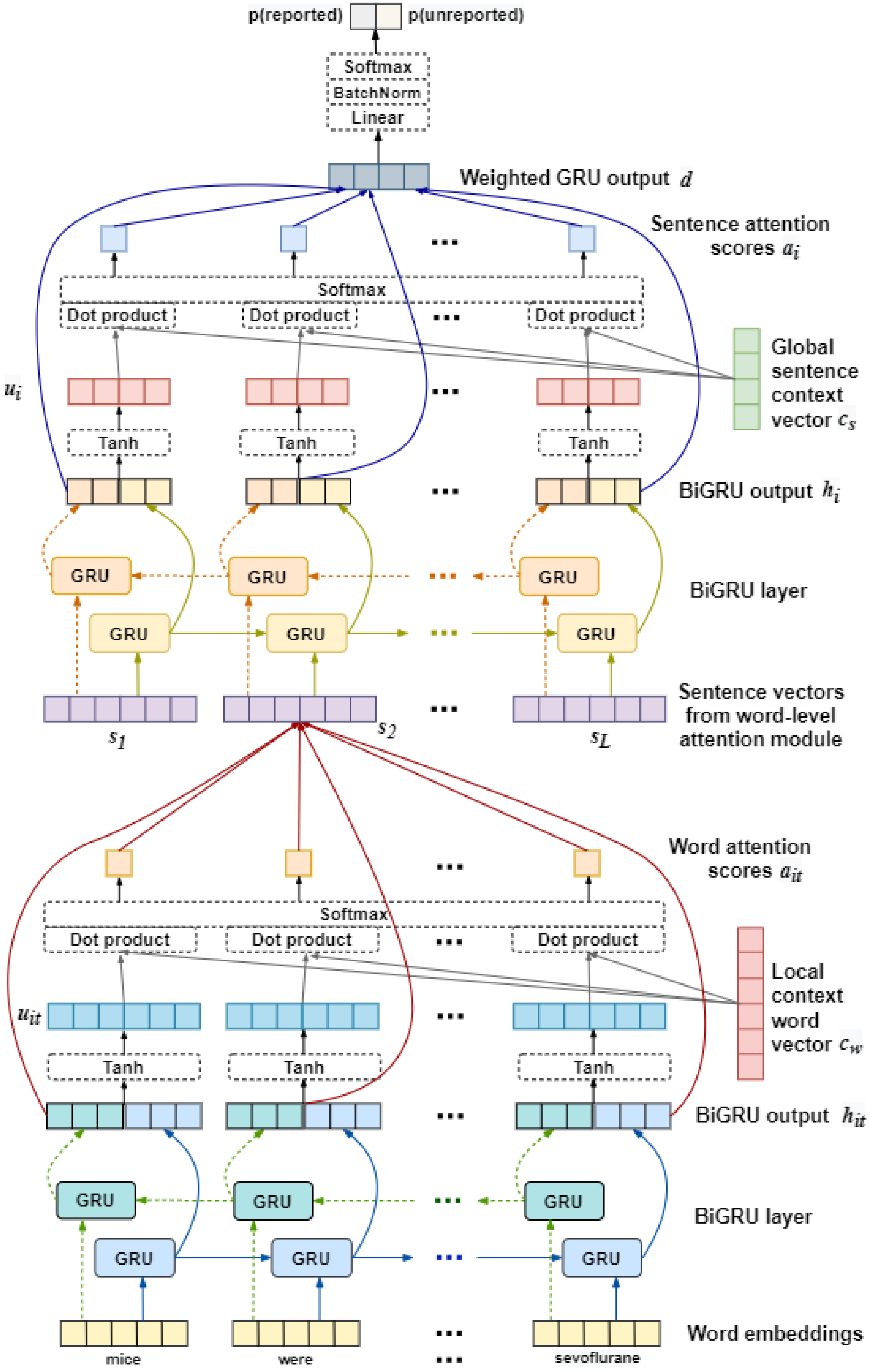

Supplementary Figure 4: The model architecture of BERT with three different head layers for long document classification. Module in red/black/blue box refers to linear head, convolutional head and LSTM head separately.

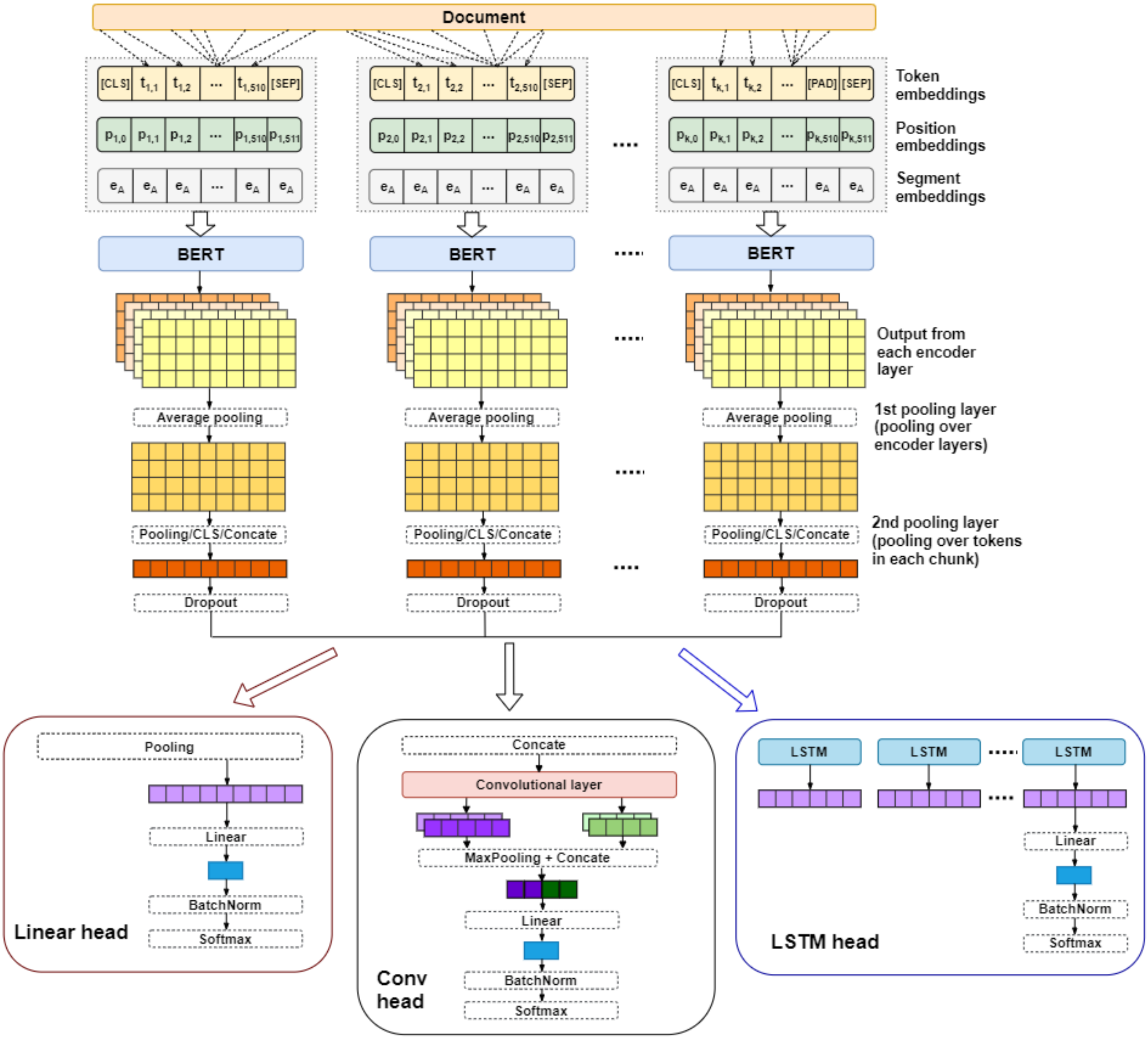
